# Mapping Gene Impact on Single-cell Transcriptomic Networks via Perturbation Response Scanning

**DOI:** 10.64898/2025.12.15.694358

**Authors:** Shreyan Gupta, Selim Romero, James J. Cai

## Abstract

Gene knockout experiments are essential for dissecting gene function, and CRISPR has made targeted gene disruption more accessible than ever. Single-cell CRISPR screening enables the construction of rich genetic perturbation landscapes, facilitating the identification of genes whose perturbation strongly reshapes cellular states. However, due to the nonlinear dependencies within gene networks, identifying the most impactful tangible genes remains challenging. Existing virtual knockout methods estimate downstream effects of single-gene deletions but do not evaluate whether such perturbations disrupt global information flow or compromise cellular robustness. To address this limitation, we adapt a perturbation-response framework originally developed for protein structural dynamics to identify gene modules most susceptible to perturbation. We introduce the single-cell Perturbation Impact Index (scPII), a data-driven metric derived from gene regulatory networks that quantifies system-level responses to gene perturbations, without using any CRISPR screening information. Our results demonstrate that scPII effectively identifies genes whose perturbation has the greatest system-wide impact. Analysis of single-cell RNA-seq data from the Cancer Cell Line Encyclopedia revealed a strong correlation between scPII-derived gene impact scores and gene effect scores from genome-wide CRISPR screens. These findings indicate that scPII provides a robust metric for quantifying gene knockout effects. More broadly, integrating perturbation response scanning with gene regulatory networks offers a powerful framework for advancing single-cell data analytics in biomedical research.

## Introduction

Understanding the architecture of gene regulatory networks (GRNs) is a central goal of systems biology, offering a roadmap for mapping complex gene functions and identifying therapeutic targets for biomedical research. A GRN is often represented as a graph with nodes as genes and edges as transcriptional (e.g., regulation, co-expression) or physical (e.g., protein-protein) links between genes [1]. Within a GRN, certain genes are critical regulators, acting as “driver nodes” or “control hubs”, on which perturbations can induce widespread transcriptional changes [2]. Pinpointing these impactful nodes is essential for disease modeling and therapeutic target discovery, but such a task remains challenging.

GRNs are often inferred using the “guilt by association” principle, which posits that functionally related genes have similar expression patterns [3]. While this is a useful starting point, resulting GRNs tend to be overly simplified. The real GRNs are inherently complex, nonlinear, and highly context-specific, dynamically adapting their regulatory landscapes in response to environmental stimuli, disease, and stress [4, 5]. Experimentally unraveling such vast combinatorial complexity [6] requires comprehensive perturbation studies such as those leveraging CRISPR-based screens [7-9], which are still insufficient owing to associated logistical challenges, survivorship bias [10], and ethical considerations [11]. Thus, computational algorithms that can accurately and efficiently model these context-specific perturbations are of critical need.

The emergence of single-cell RNA sequencing (scRNA-seq) has enabled the reconstruction of GRNs with an unprecedented high resolution [12]. However, accurately identifying critical genes remains a key challenge because: (1) inferred GRNs from scRNA-seq data need to be validated for their robustness, and (2) an efficient algorithm is needed to quantify a gene’s network-wide impact to identify impactful nodes. The first problem has been largely solved, given that GRN construction from scRNA-seq data is an active area of research [13-15]. We have systematically evaluated several algorithms and confirmed that GRNBoost2 has demonstrated superior performance in both overall inference and hub gene identification [16]. For the second problem, it is crucial to use a computationally efficient approach to quantify the global impact of a gene, taking into account both direct and indirect interactions [17]. Existing methods like scTenifoldKnk [18] and GenKI [19] can simulate single gene knockouts (KO) and quantify their effect on other genes, but to get a global impact, these algorithms have to iterate over every gene in a GRN, which is extremely computationally intensive. Thus, their utility in estimating the overall gene impact landscape has not been feasibly exploited.

Network diffusion, a process by which the effect of perturbation spreads through the network [20, 21], is a viable option. Especially, perturbation response scanning (PRS), a method for studying conformational changes undergone by proteins under selected external perturbations [22]. PRS relies on systematically applying forces at singly selected residues and recording the linear response of the whole protein. The protein in the folded state is equivalent to a three-dimensional elastic network. Similarly, the PRS approach has been adapted to identify critical genes and repurposing drugs using curated GRNs by simulating the spread of a perturbation through the network.

In this study, we adapt the PRS approach and apply it to weighted single-cell GRNs, using the graph Laplacian to provide a computationally efficient and mathematically rigorous solution for identifying impactful nodes. We term the *single-cell perturbation impact index* (scPII) as a measure of the extent to which the perturbation of one gene globally impacts the entire regulatory network. Our algorithm uses GRNBoost2 to construct a robust GRN and then applies PRS to compute the index, providing a scalable, data-driven solution for identifying critical genes. Compared to existing virtual KO solutions, scPII is a more computationally efficient method, providing significant biological insights in our case studies. First, we validate scPII by applying it to a curated GRN from BioGRID [23] to identify key points of perturbations and drug targets in human glioblastoma. We then systematically benchmark scPII against several off-the-shelf network centrality metrics and two dedicated virtual KO methods (scTenifoldKnk [18] and GenKI [19]) using 16 pan-cancer scRNA-seq datasets from the Cancer Cell Line Encyclopedia (CCLE) [24] and real-world CRISPR KO screens from the DepMap portal [25] and BioGRID-ORCS database [26].

## Results

### The scPII framework

The scPII framework is developed based on the PRS method, which was originally developed for analyzing tertiary protein structures [22] and later adapted for gene network analysis [27]. The scPII score quantifies the impact of individual gene perturbations on GRNs derived from scRNA-seq data. The computing process of scPII is depicted in Figure 1 and detailed below.

**Figure 1.**
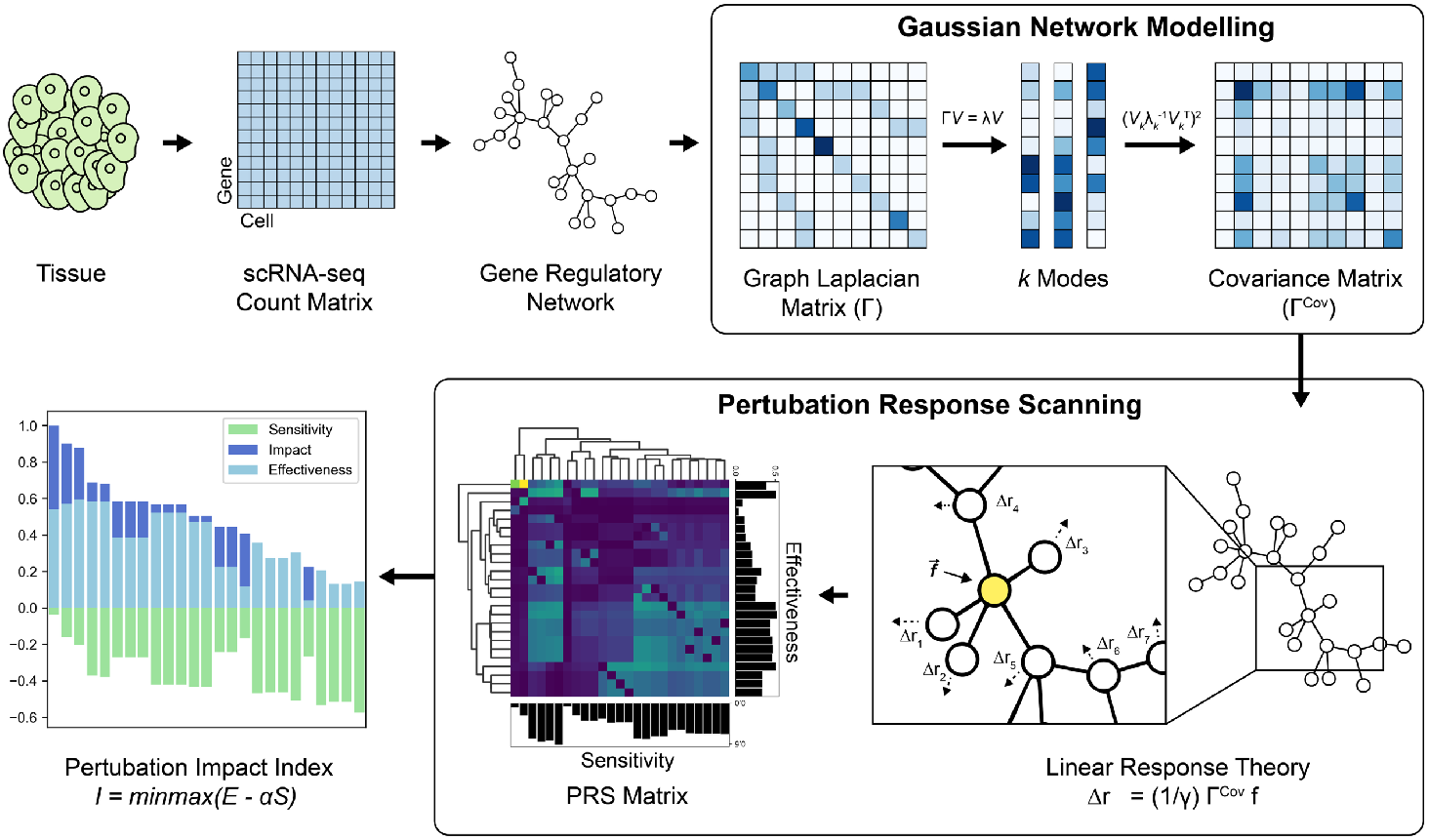
Overview of single-cell perturbation impact index (scPII) computation workflow. The scPII algorithm begins by constructing a GRN from scRNA-seq data. This GRN is regarded as a Gaussian network model, and its graph Laplacian matrix is calculated. The eigen-decomposition of the graph Laplacian is used to identify the slowest modes of the network, which capture cooperative behaviors. These slowest modes are then utilized to construct a covariance matrix (Γ^Cov^). This covariance matrix is subsequently normalized by its diagonal elements to generate a PRS matrix. Finally, effectiveness and sensitivity scores are derived from the PRS matrix, which are then integrated to compute each gene’s impact index.

We begin with the construction of an undirected GRN using scRNA-seq data. The largest connected component of the resulting GRN is retained for analysis in the form of a graph adjacency matrix, which is then used to compute the graph Laplacian matrix, Γ. The Laplacian matrix serves as the foundation for modeling network dynamics. In this analytical framework, the GRN is considered as a Gaussian network model, conceptualizing the network structure as nodes connected by springs. Such a model facilitates the analysis of collective motions and inter-node interaction topology. Key to this approach is the computation of an approximate pseudo-inverse of the Laplacian, Γ^+^, which captures cross-correlation interactions and node relationship fluctuations. This pseudo-inverse is efficiently estimated by the lowest *k* non-zero eigenvalues and their corresponding eigenvectors from the eigen-decomposition spectra of Γ. The squared elements of Γ^+^then form the covariance matrix, Γ^Cov^, which maps the response of other nodes when a target node is perturbed.

The PRS method is applied by simulating a unit force on each node and estimating the resulting displacement of all other nodes, analogous to linear response theory [28]. The scaled PRS matrix is then constructed, where each row represents the influence profile of a perturbed node. We derive two fundamental node-specific metrics using the row-wise and column-wise means of the PRS matrix: effectiveness (𝔼) and sensitivity (𝕊), respectively. 𝔼 quantifies a node’s ability to propagate perturbation information throughout the network. Conversely, 𝕊 measures a node’s susceptibility to external perturbations from its neighbors. Both scores are log-transformed to mitigate outlier effects.

Finally, the impact index (or scPII score) is computed as 𝕀 = 𝔼 − *α*𝕊, a scaled difference between 𝔼 and 𝕊. The index provides a balanced quantification of a gene’s overall importance within the network. It reflects each gene’s capacity to induce structural changes in the GRN while accounting for its inherent susceptibility to external influences.

Genes with greater values of scPII score are those that effectively perturb the network while being less prone to external influences, theoretically making them prime candidates for targeted therapeutics.

### ScPII identifies drug targets in glioblastoma networks

We first tested whether scPII can identify genes with high impact on the network structure using an established, human-curated GRN. We obtained the curated GRN derived from the Glioblastoma Multiforme (GBM) project from the BioGRID database [23]. As detailed in **Methods**, nodes (n = 10,924) in the network represent genes and edges (n = 32,958) represent the experimentally validated interactions between genes. For functional validation, we used a comprehensive set of CRISPR screens targeting GBM cell lines, retrieved from the DepMap and the BioGRID-ORCS databases [29] (**Fig. 2a**). With these datasets, we performed a direct comparison of the computationally inferred perturbation impact with experimentally observed phenotypic consequences.

**Figure 2.**
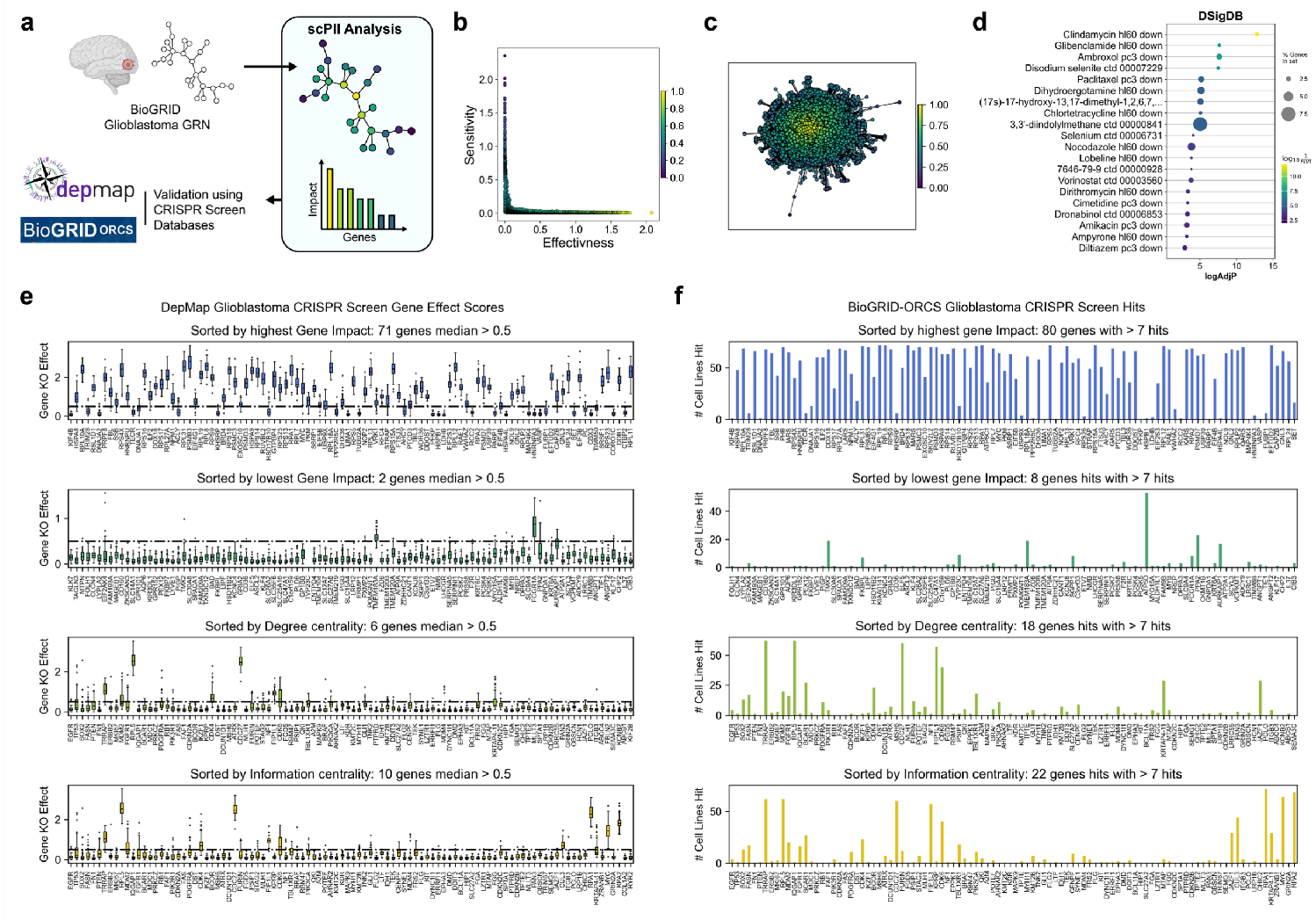
scPII identifies highly impactful genes in glioblastoma multiforme (GBM) networks and predicts drug targets. **(a)** Schematic overview of the scPII application workflow. A curated glioblastoma GRN from BioGRID is fed into the scPII analysis framework to compute each gene’s perturbation impact. The predicted top genes are then validated using their perturbation profile information from the real CRISPR gene KO screens in the DepMap and BioGRID-ORCS databases. **(b)** Force-directed graph representation of a GBM GRN, where node color intensity corresponds to the value of the scPII score. **(c)** Scatter plot illustrating the relationship between gene sensitivity, effectiveness, and scPII score. Each point represents a gene. **(d)**Drug enrichment analysis results for the top 100 scPII-identified genes, using Enrichr against the DSigDB database. Drugs are sorted by the increasing adjusted P values. **(e)** Distributions of gene KO effect scores for the top 100 genes, sorted by different centrality metrics, against the negative control of low-impact genes. The top panel shows genes sorted by the highest scPII value. The second panel shows genes sorted by the lowest scPII value. The third panel shows genes sorted by degree centrality. The bottom panel shows genes sorted by information flow centrality. The dashed line at 0.5 indicates the threshold for a functionally significant gene KO effect. **(f)** The number of GBM cell lines significantly affected (hit) by CRISPR-based KO of the top 100 genes, sorted by different centrality metrics, against a negative control of low-impact genes. The top panel shows genes sorted by the highest scPII value. The second panel shows genes sorted by the lowest scPII value. The third panel shows genes sorted by degree centrality. The bottom panel shows genes sorted by Information flow centrality.

The fully connected GBM GRN was fed into the scPII framework. For each gene, three key metrics: effectiveness (𝔼), sensitivity (𝕊), and impact index (𝕀), were calculated (**Supplementary Table 1**). As mentioned, 𝔼 measures the level to which a gene’s perturbation alters network topology, 𝕊 the level of the gene’s susceptibility to external perturbations. When each gene’s impact was visualized on a force-directed graph, we observed that centrally located genes generally exhibited higher impact values, while genes located peripherally exhibited lower impact values (**Fig. 2b**). This relationship is consistent with our theoretical assumptions, and sensitivity was observed to be largely inversely proportional to its effectiveness. Consequently, genes with high impact scores typically demonstrate high effectiveness and low sensitivity (**Fig. 2c**).

To explore the clinical implications of the genes prioritized by scPII, we performed an enrichment analysis on the top 100 scPII-identified genes using Enrichr [30] and the DsigDB [31] database—a comprehensive repository of drugs and their known gene targets (**Fig. 2d, Supplementary Table 2**). Our results revealed a significant enrichment for several drugs with established or emerging relevance in GBM therapy. Clindamycin [32], the top enriched drug, has previously demonstrated cytotoxic effects on GBM tumors. Sodium selenite [33] and selenium [34] were also identified, both promising anti-GBM agents. Paclitaxel [35, 36], the well-known broad utility chemotherapeutic agent in cancer treatment, was enriched. Chlortetracycline [37] was found to be enriched, a drug known to inhibit the ADP-ribosylation factor subfamily of the RAS superfamily, thereby suppressing the invasive capacity of glioma. Nocodazole [38], an established agent inducing transient mitotic arrest, has been investigated in glioma cell lines for therapeutic purposes was identified as well. Lobeline was also enriched, a drug shown to induce GBM cell death through delayed hypoxic stress recovery. GBM cells have demonstrated enhanced sensitivity to dirithromycin [39]. Cimetidine [40, 41] was also identified, known for its anti-cancer effects, particularly in combination therapies for GBM. Furthermore, several other enriched drugs, including glibenclamide [42], ambroxol [43], and dihydroergotamine [44], have exhibited anti-cancer effects in various combination therapies, suggesting potential broader utility, including in GBM. Notably, epidemiological studies have indicated a lower incidence of glioblastoma multiforme in patients receiving the ion channel blocker diltiazem [45]. The direction of the gene sets was also down for most pathways, signifying that these pathways contained genes that were downregulated by the drug.

### Comparing scPII with network centralities

In network theory, centralities are used to quantify the relative importance or influence of nodes within a network. Here, to benchmark scPII’s performance, we compared scPII against two network centrality measures: degree centrality and information flow centrality. The latter is particularly relevant for comparison as it, like scPII, utilizes the inverse of the graph Laplacian matrix for its computation. Because of this shared mathematical foundation, we chose to test its performance on a real biological network.

For the benchmark, we obtained data, consisting of 18,443 genes in 50 CRISPR screens of GBM-derived cell lines, from the DepMap portal. After computing the impact and the two centrality measures, we selected the top 100 genes according to each metric (**Supplementary Table 3**). These gene sets were then compared to their corresponding gene KO effect scores, obtained from real CRISPR screens in the DepMap portal (**Fig. 2e**). We observed that 71 genes identified by the scPII metric had a median absolute gene KO score greater than 0.5. In contrast, only 6 genes sorted by degree centrality and 10 genes sorted by information centrality met this threshold. This result indicates that our impact index has a higher predictive power. As a crucial negative control, genes associated with the lowest impact scores were also examined; only 2 of these exhibited a median gene KO effect score greater than 0.5, further validating the specificity of scPII.

We further validated the genes using GBM-associated CRISPR screens from the BioGRID-ORCS database (**Fig. 2f**). A total of 72 screens were obtained, consisting of 18,888 unique gene perturbations. Each gene’s perturbation was characterized as a ‘hit’ or a ‘miss’, with a hit meaning a significant effect on cell proliferation of the cell line. In our tests, we consider a gene to be a successfully predicted perturbation candidate if it significantly affects (hits) at least 10% (7) of the 72 cell lines. Among the top 100 genes sorted by impact, 80 genes had hits on more than 7 cell lines. Meanwhile, when sorted by Degree centrality and information flow centrality, only 18 and 22 genes, respectively, had hits in more than 7 cell lines. As a negative control, the last 100 genes, when sorted by impact, only 8 genes had hits in more than 7 cell lines.

In summary, the scPII framework consistently identified high-impact genes that significantly influence the overall survivability and proliferation of GBM cell lines from real-world datasets. Moreover, the genes predicted by scPII represent potential targets for existing drugs with known efficacy against GBM. These findings stress scPII’s potential as an effective tool for high-throughput drug screening and repurposing efforts in cancer therapy. In the next section, we apply scPII to real single-cell data-driven GRNs.

### ScPII predictions are consistent with large-scale CRISPR knockout data

Next, we applied the scPII framework to GRNs derived from scRNA-seq datasets. We obtained scRNA-seq data from the cancer cell line encyclopedia (CCLE) database, alongside paired CRISPR KO screens from the DepMap portal for robust perturbation response validation. To ensure data quality and sufficient cellular resolution, we only retained CCLE cell lines with over 500 cells, resulting in 16 suitable datasets for analysis (**Fig. 3a**), across 9 different tissue types (**Fig. 3b, Supplementary Table 4**).

**Figure 3.**
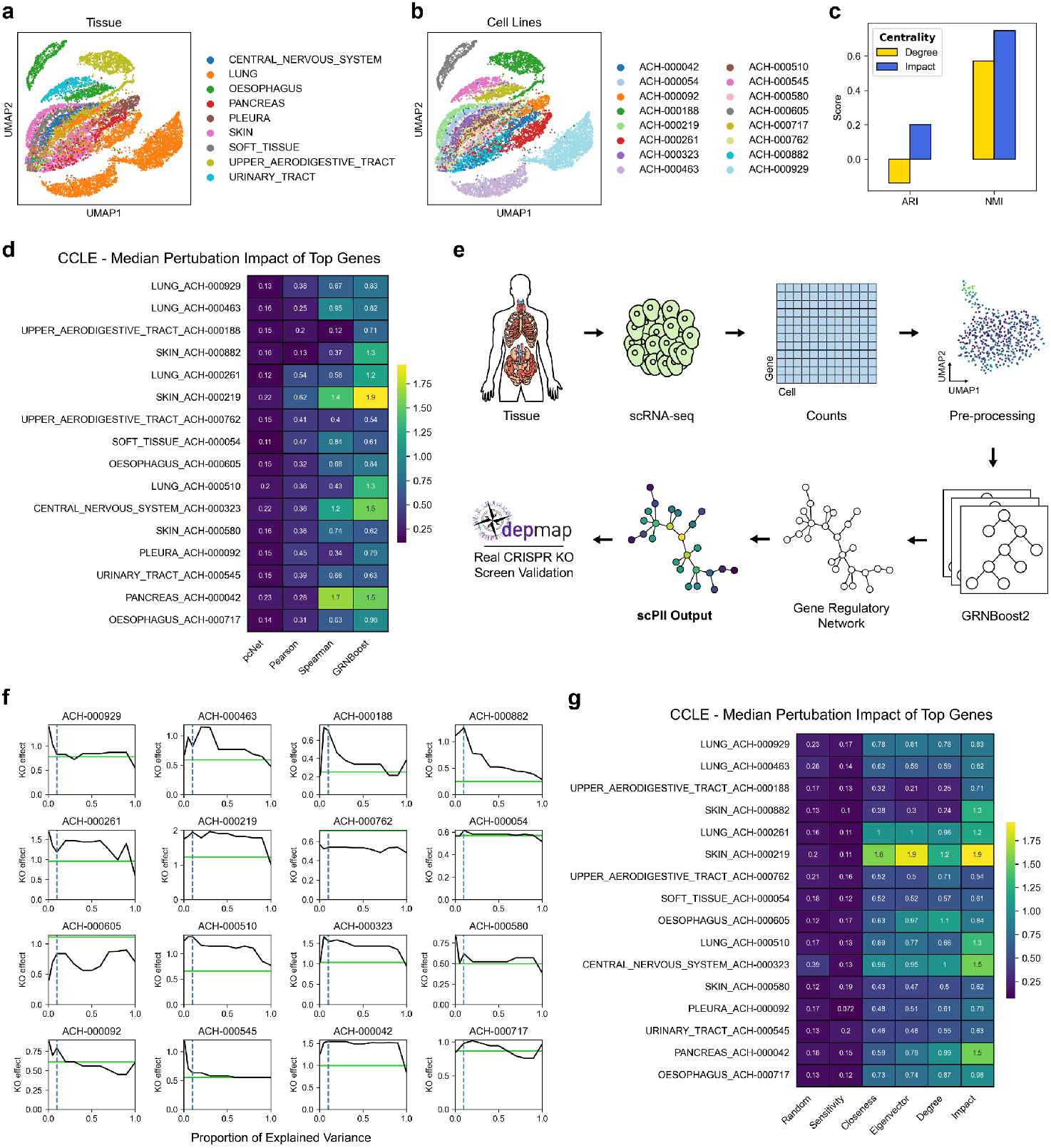
scPII identifies highly impactful genes in data-driven networks from 16 cancer cell lines and predicts KO target genes. **(a, b)** UMAP plots visualizing the clustering of 16 CCLE cancer cell lines colored by **(a)** different tissue types and **(b)** different cell lines. **(c)** Bar plot comparing the median Adjusted Rand Index (ARI) and Normalized Mutual Information (NMI) scores for clustering 16 CCLE cancer cell lines using scPII Impact indices versus Degree centralities of their common genes. **(d)** Heatmap showing the median gene KO effect scores for the top 50 genes ranked by four different GRN inference methods (Pearson, Spearman, pcNet, and GRNBoost2) across 16 CCLE cell lines. The color intensity and annotated values indicate the median KO effect score. **(e)** The scPII pipeline from raw scRNA-seq data to GRN inference and final scPII output. The diagram shows the following steps: 1) tissue extractions, 2) single-cell isolation and sequencing, 3) mapping and generation of counts, 4) preprocessing, 5) GRN construction using GRNBoost2, and 6) scPII analysis, which was finally validated against real DepMap CRISPR KO screens. **(f)** Median gene KO effect scores for varying proportions of explained variance in the scPII algorithm across eight representative cell lines. The x-axis represents the proportion of explained variance, and the y-axis represents the median KO effect. The dashed lines highlight the optimal 0.1 (10%) explained variance threshold. **(g)** Heatmap comparing the median gene KO effect scores of the top 50 genes ranked by four different centrality measures (random, closeness, eigenvector, degree) and the scPII Impact score across the 16 CCLE cell lines.

The first step of the scPII framework is the construction of a reliable data-driven GRN from the scRNA-seq count data. To find the ideal candidate GRN construction algorithm, we benchmarked four distinct methods: Pearson correlation, Spearman correlation, pcNet, and GRNBoost2, as detailed in the Methods section. These GRNs served as the input for the scPII algorithm, which then predicted the Impact of each gene within each cell line. To validate the predictive power of scPII using each GRN construction tool, for every cell line, we extracted the top 50 genes, ranked by their scPII Impact score, and compared them with their absolute gene KO effect scores obtained from the paired DepMap CRISPR screens (**Supplementary Fig. 1a**).

Across the 16 analyzed cell lines, the combination of GRNBoost2 for GRN construction paired with scPII consistently yielded the highest median gene KO effect scores for the top 50 genes in 11 of the 16 cases (**Fig. 3d**). Thus, we selected GRNBoost2 as the optimal GRN inference method for use in the scPII framework (**Fig. 3e**). Spearman correlation was the next best performer.

The covariance matrix, Γ^*Cov*^, computation in the scPII framework is contingent on selecting the eigen vectors associated with the lowest *k* eigen values. Given the varied number of genes across the diverse scRNA-seq datasets used in this study, a fixed *k* value would be suboptimal. Instead, we dynamically tested the optimal *k* by selecting the eigenvalue-vector pairs that collectively explained a specific proportion of the total variance captured during eigen decomposition. We systematically evaluated different variance proportions (**Fig. 3f**) and found that 0.1 (10%) of explained variance consistently corresponded to the highest median gene KO effect in real CRISPR KO screens. This optimized parameter ensures robust performance across datasets with varying network sizes.

Now with GRNBoost2 as the optimal GRN construction algorithm, we tested scPII’s performance against three established graph centrality measures: closeness, eigenvector, and degree centrality. For each of the 16 scRNA-seq-derived GRNs, we computed these centrality measures along with the impact index from scPII. For the top 50 genes ranked by each centrality measure, along with 50 randomly selected, we recorded their median absolute gene KO effect scores from their corresponding DepMap CRISPR screens (**Fig. 3g, Supplementary Fig. 1b**). scPII consistently outperformed the conventional centrality measures, yielding the highest median gene KO effect scores in 14 out of the 16 cases. Conversely, as a critical negative control, we examined the top 50 genes with the lowest impact scores. These low-impact genes consistently exhibited the lowest median gene KO effect in 13 out of the 16 cases, further validating the predictive specificity of scPII’s impact index.

We hypothesized that similar cell lines should have similar perturbation response patterns and thus they should cluster together based on the impact indices of their genes. Hence, to assess the stability of information across datasets by scPII, we compared the clustering of the 16 CCLE cancer cell line GRNs using scPII Impact indices versus Degree centralities of their common genes. The impact indices and degree centralities for these shared genes were stored in a matrix, and the cell lines were clustered (**Supplementary Fig. 2**). We then evaluated these clustering results against the original tissue sources of the cell lines. The impact index yielded higher adjusted Rand index (ARI) and normalized mutual information (NMI) scores compared to degree centrality (**Fig. 3c**). This suggests that cancer cell lines, when clustered by the impact indices of their genes, similar cell lines tend to have similar impact metrics irrespective of the number of connections of each node.

These results demonstrate that scPII, when optimally paired with GRNBoost2 for data-driven network inference, robustly identifies functionally impactful genes from scRNA-seq data that were experimentally validated against real CRISPR KO screens. In the next section, we proceed to test the efficiency of scPII against established virtual KO algorithms.

### Benchmarking scPII against established virtual KO frameworks

Given the large-scale, high-dimensional data structures in the field of transcriptomics, the practicality and scalability of computational tools are just as important as their accuracy. Hence, in this section, we performed a comparative analysis (**Fig. 4a**) of our proposed scPII framework against established virtual KO methods, including scTenifoldKnk and GenKI, using GRNs from three distinct CCLE datasets: ACH-000463, ACH-000882, and ACH-000762. For both virtual KO methods, each gene in the network for systematically knocked out to compute response scores as detailed in the methods. Each network comprised over 3,300 genes. We also included Degree centrality as a baseline method derived from intrinsic network-based properties. The performance of each method was validated using corresponding CRISPR KO screen data from the DepMap portal.

**Figure 4.**
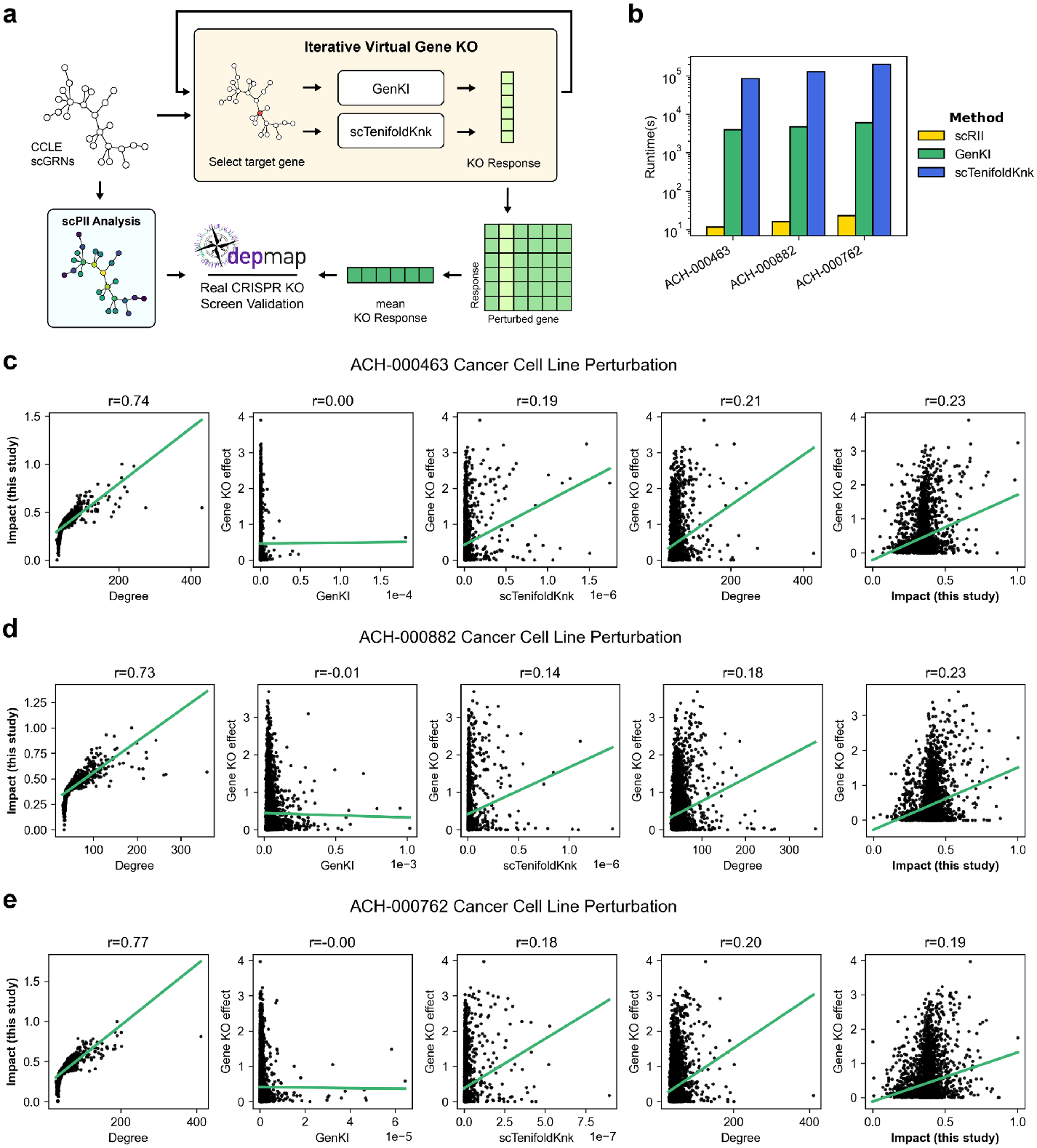
Comparative analysis of scPII with virtual knockout simulation methods. **(a)** Overview of the comparative framework. GRNs inferred from CCLE scRNA-seq data are subjected to iterative virtual gene KO using either GenKI or scTenifoldKnk. The resulting KO responses are then averaged to generate a mean KO response score. These scores, along with scPII Impact scores, are validated against real CRISPR KO screen data from the DepMap portal. **(b)** Runtimes of scPII, GenKI, and scTenifoldKnk across three representative CCLE cell lines (ACH-000463, ACH-000882, ACH-000762). **(c, d, e)** Scatter plots illustrating the Pearson correlation (r) between impact, response scores from Virtual KO methods, and real CRISPR gene KO effect scores for **(c)** ACH-000463, **(d)** ACH-000882, and **(e)** ACH-000762 cancer cell lines, respectively.

We first assessed the computational efficiency of all three algorithms (**Fig. 4b**). scPII consistently demonstrated significantly faster runtimes compared to both scTenifoldKnk and GenKI. For all three datasets, scPII’s runtime was as little as three orders of magnitude of that of scTenifoldKnk or GenKI (**Supplementary Table 5**).

Next, we evaluated the accuracy of KO impact prediction by calculating the Pearson correlation coefficient (r) between each algorithm’s output and the experimental CRISPR KO data. For ACH-000463 and ACH-000882 (**Supplementary Tables 6 and 7**), scPII exhibited the highest correlation (r = 0.23, in both cases, **Fig. 4c,d**), which was substantially higher than the correlations observed for GenKI (r = −0.01 and r = 0, respectively) and for scTenifoldKnk (r = 0.19 and r = 0.14, respectively). Degree centrality showed a strong correlation (r = 0.21 and r = 0.18, respectively), but both were lower than that of scPII.

For ACH-000762 (**Fig. 4e**), the results were slightly different (**Supplementary Table 8**). In this cell line, degree centrality showed the highest correlation with CRISPR KO effects (r = 0.20), followed closely by scPII (r = 0.19). Both methods significantly outperformed GenKI (r = 0.00) and scTenifoldKnk (r = 0.18). These findings align with our earlier observation that genes prioritized by degree in this specific cell line had the highest median CRISPR KO effect scores (**Fig. 3g**). The strong mutual correlation between scPII and degree centrality (such as r = 0.77 for ACH-000762) highlights a statistical overlap in how these two metrics identify high-impact genes in GRNs. Nevertheless, scPII outperforms degree centrality as it does not prioritize hub genes.

Thus, we demonstrated superior performance of scPII in predicting real KO effects by offering a significant computational efficiency advantage over existing virtual KO methods. Our results suggest that scPII is an effective and scalable tool for identifying genes based on their impact on a cellular network.

## Discussion

The PRS approach has been adapted previously in analyzing genetic interaction networks to identify effector genes (high effectiveness in distributing information) and sensor genes (high sensitivity in receiving information) [27]. Inspired by this, we extended the principles of PRS to GRNs derived from scRNA-seq experiments. In this study, we introduced the single-cell perturbation impact index (scPII) to efficiently quantify the impact of genes within dynamic, data-driven single-cell GRNs. The scPII method provides a powerful and scalable solution for identifying critical regulatory genes, or “control hubs”, whose perturbation can have a widespread effect on the cellular regulatory landscape.

To validate scPII, we first applied it to a curated GRN from BioGRID, where it successfully identified genes with significant real-world KO effects from CRISPR screens on the DepMap portal. scPII consistently outperformed established centrality measures in this controlled setting. As a compound index, scPII combines perturbation effectiveness (the ability to propagate perturbations) and sensitivity (susceptibility to external forces), proving to be a biologically relevant measure of a gene’s perturbation impact on the system. The inverse relationship we observed between effectiveness and sensitivity for high-impact genes suggests that, for most genes, if they are powerful diffusers of information, then they tend to be robust to perturbation. The above finding is biologically compelling as it aligns with the concept of “control hubs”, i.e., these high-impact genes are not merely well-connected but are topologically positioned to exert control while remaining resilient to external network noise.

Beyond identifying high-impact genes, our analysis of the GBM network showed that these genes are significantly enriched for targets of drugs with established efficacy against GBM. The recovery of compounds such as clindamycin, paclitaxel, and sodium selenite not only validates scPII’s biological relevance but also highlights its utility for high-throughput drug repurposing and target prioritization. By computationally predicting genes associated with clinically relevant therapeutics, scPII offers a powerful translational tool capable of accelerating discovery pipelines. Notably, while this manuscript was being prepared, a new study [46] reported the successful application of the PRS method for drug repurposing, underscoring the growing importance of perturbation-based network approaches in therapeutic discovery..

We also applied our framework to 16 unique cancer cell lines from the CCLE database. This pan-cancer analysis further solidified scPII’s utility. The genes ranked highest by scPII consistently showed higher median KO effect scores in real DepMap CRISPR KO screens than genes ranked by conventional centrality measures or existing virtual KO methods. This demonstrates scPII’s robustness and predictive power, especially when handling the inherent noise and dynamic nature of real-world data-driven scRNA-seq GRNs. Moreover, despite significant differences in the gene numbers and topology of the GRNs derived from each dataset, the Impact scores of common genes were remarkably conserved across similar cancer cell lines. In contrast, graph centrality-based measures like Degree centrality failed to maintain this information across the cell lines.

After testing several GRN construction methods, we found that GRNBoost2 was the most effective for our scPII workflow. This aligns with findings from Stock et al. [13], which noted GRNBoost2’s superior ability to identify hub genes. Our analysis showed that GRNBoost2 produced higher predictive performance than the Pearson’s correlation-based GRNs used by Acar et al. [27] in past studies.

To improve the scPII’s performance further, we recommend using a weighted Laplacian for the PRS, which avoids the arbitrary selection of a Pearson correlation cutoff (e.g., 0.2). This approach prevents significant changes to network topology that can occur with cutoff-based methods [47], ultimately providing more robust PRS results. Based on our findings, we suggest using GRNBoost2 for GRN construction and a weighted Laplacian for the PRS to enhance scPII performance. However, we acknowledge and stress that the most suitable algorithm for GRN construction may vary depending on the specific sample and context.

We developed a more robust approach for eigenvalue selection during covariance matrix construction, which makes our analysis more adaptable to diverse datasets with varying GRN sizes and structures. Previous methods relied on an arbitrary number of 20 principal components [48], which is not context-specific and can drastically alter the global information preserved. To overcome this, we introduce a dynamic, data-driven approach that uses a variance-based cutoff to ensure scPII outputs are more robust and translatable. We recommend capturing 10% of the total variance during the covariance matrix computation for GNM analysis, as this was rigorously validated to provide ideal predictive performance.

Furthermore, scPII offers a significant advantage in computational efficiency. Our comparative analysis against virtual KO simulation methods, such as scTenifoldKnk and GenKI, shows that scPII operates at least two orders of magnitude faster. While scTenifoldKnk and GenKI are effective for simulating individual gene KOs, their iterative nature makes them computationally intensive for ranking the impact of every gene in large, high-dimensional datasets. scPII’s algebra-driven framework bypasses this bottleneck, making it a highly scalable tool for large-scale single-cell transcriptomic data analysis.

A current limitation of scPII is that the impact metric only quantifies the magnitude of perturbation effect on the GRN (transcriptional landscape), but it does not predict its direction (e.g., whether it will promote proliferation or induce cell death). For this reason, we utilized the absolute KO effect scores from the DepMap portal for all our testing scenarios. Therefore, the impact index serves as a measure of overall impact, not a predictor of phenotypic outcome. A possible future direction will be to focus on predicting the directionality of perturbation impact to further enhance scPII’s predictive capabilities. Additionally, while this study focused on the impact of perturbing individual nodes, exploring the efficacy of perturbing network edges is a crucial direction for future research [2].

In conclusion, scPII represents a significant advance in computational genomics. Its ability to accurately and efficiently identify functionally impactful genes, predict drug targets, and provide a stable measure of gene importance across diverse cellular contexts addresses a long-standing need in the field. As scRNA-seq datasets continue to grow, tools like scPII will be essential for translating these vast datasets into actionable biological insights and therapeutic strategies.

## Methods

### Acquisition of the BioGRID GBM dataset

For our analyses, we obtained a curated human GRN from the BioGRID project portal (https://thebiogrid.org/) [23]. This network was specifically derived from Glioblastoma Multiforme (GBM) tumor samples from The Cancer Genome Atlas (TCGA) and consists of 10,924 unique genes (nodes) and 32,958 regulatory interactions (edges).

### Acquisition and pre-processing of scRNA-seq data sets

We obtained publicly available scRNA-seq datasets of cancer cell lines from the Cancer Cell Line Encyclopedia (CCLE) project. Raw scRNA-seq data of ∼200 cancer cell lines were obtained from the Broad Institute’s single-cell portal (SCP542) [24].

We implemented a standard scRNA-seq data analysis pipeline using the Scanpy (v1.9.8) Python package [49] for data preprocessing. We excluded cells if they had fewer than 1,000 total counts, fewer than 500 unique expressed genes, or greater than 5% mitochondrial gene expression. Genes expressed in less than 20% of the remaining cells were also removed. We then normalized the count data using the ‘*normalize_total*’ function and then *log+1* transformed it using the ‘*log1p’* function. Next, we scaled and centered the data for principal component analysis (PCA). Uniform manifold approximation and projection (UMAP) was performed using the top 10 principal components to generate a 2D representation of the cells. The log-normalized count data were used for GRN construction.

To ensure the availability of functional validation data, we excluded cell lines lacking corresponding CRISPR-Cas9 KO screens in the DepMap portal [29] from further analysis. Following all preprocessing and filtering steps, only cell lines with a minimum of 500 cells were retained to ensure sufficient cellular resolution and minimize potential errors during the GRN construction phase. This was done to preserve enough cells to minimize error during the GRN construction step. This yielded 16 candidate cell lines for subsequent experimental validation and analysis (**Supplementary Table 3**).

### Acquisition of real CRISPR gene KO screens from the DepMap portal

To validate the computationally inferred gene impacts, we acquired genome-wide CRISPR-Cas9 gene KO screen data from the DepMap portal (https://depmap.org/portal/) [29]. This comprehensive dataset provides quantitative gene effect scores, which reflect the impact of individual gene perturbations on cell viability across a large panel of cancer cell lines. For each CCLE cell line and BioGRID dataset included in our analysis, the corresponding absolute gene KO effect scores were extracted and utilized as the experimental ground truth for evaluating the predictive performance of our algorithms.

### Acquisition of real CRISPR gene KO screens from the BioGRID-ORCS database

We used genome-wide CRISPR screen data from the BioGRID-ORCS database v2.0.18 (https://orcs.thebiogrid.org/) [26]. We only selected screens that measured changes in overall cell proliferation. This resulted in a dataset of CRISPR screens from 72 different cell lines, all associated with GBM. In each screen, a gene was recorded as a “hit” (a binary feature) if its perturbation had a significant effect on the cell line.

### Gene regulatory network construction

#### Network construction by correlation

We used two correlation metrics to construct GRNs: Pearson and Spearman. Pearson correlation coefficient, a widely used measure of linear association between gene expression profiles. For a given cell type, the gene expression count matrix was represented as *X* ∈ ℝ^*p*×*n*^, with *p* genes and *n* cells. The gene expression level of the *i*^*th*^ gene in all cells is represented by the *i*^*th*^ row of *X*, denoted by *X*_*i*_ ∈ ℝ^*n*^. We calculated the Pearson correlation coefficient, *r*, and the rank-based Spearman correlation coefficient, *ρ*, for each pair of genes, *X*_*i*_ and *X*_*j*_, and stored the coefficient in the correlation matrix, *A*.

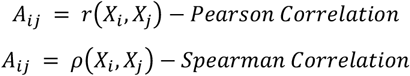

The correlation matrix then acts as the weighted adjacency matrix of the intracellular gene correlation network, with all diagonal entries set to 0 to remove self-loops. Entries with a correlation coefficient less than 0.1 were removed to exclude less significant interactions.

#### Network construction by PC regression

We adopted a PC regression framework, pcNet, specifically tuned for scRNA-seq data [50, 51]. For a given cell type, the gene expression count matrix is represented as *X* ∈ ℝ^*p*×*n*^ with *p* genes and *n* cells. The gene expression level of the *i*^*th*^ gene in all cells is represented by the *i*^*th*^ row of *X*, denoted by *X*_−*i*_. *X*_*i*_ denotes the matrix by deleting *X*_*i*_ from *X*. We constructed the PC regression model (for the matrix *X*_−*i*_) to estimate the effects of other *p* − 1 genes on the *i*^*th*^ gene. First, we applied principal component analysis (PCA) to 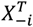 and selected the first *M* leading PCs to construct 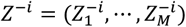, where 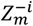 is the *m*^*th*^ PC of 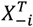, *m* = 1,2, …, *M* and *M* ≪ *min* (*p, n*). The PC loading matrix is denoted by *V*^*i*^, where *V*^*i*^ satisfies (*V*^*i*^)^*T*^ *V*^*i*^ = *I*_*M*_. We then computed the regression coefficients by regressing *X*_*i*_ on *Z*^−*i*^ using the ordinary least squares (OLS) method:

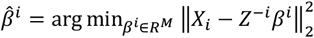

Then, the effects of the other *p* − 1 genes on the *i*^*th*^ gene was obtained by 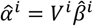. We repeated this process *p* times, with one different gene as the response gene each time to generate the *p* × *p* set of gene effects, 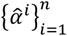, which then acts as the weighted adjacency matrix, *A*, of the intracellular gene regression network with all diagonal entries set to 0 to remove self-loops. The adjacency matrix was symmetrized and made non-negative by calculating 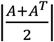. Entries less than 0.1 were removed to exclude less significant interactions.

#### Network construction by GRNBoost2

GRNBoost2 utilizes a gradient boosting machine (GBM) algorithm, specifically the LightGBM implementation, to model the regulatory relationships [16]. For a given cell type, the gene expression count matrix is represented as *X* ∈ ℝ^*p*×*n*^ with *p* genes and *n* cells. For each target gene, *X*_*i*_, GRNBoost2 treats the expression levels of all other *p* − 1 genes as potential predictors. The GBM algorithm iteratively builds an ensemble of decision trees to predict the expression of *X*_*i*_. We repeated this process for each of the *n* genes, treating each gene as a target gene in turn. This yielded an *p* × *p* matrix, *A*, where *A*_*ij*_ represents the importance score of gene *j* in predicting the expression of gene *i*. The resulting matrix, *A*, serves as the weighted adjacency matrix of the intracellular gene regulatory network. We set all diagonal entries of *A* to 0 to remove self-loops. The adjacency matrix was symmetrized and made non-negative by calculating |*max*(*A, A*^*T*^)|. Entries less than 0.1 were removed to exclude less significant interactions.

### Perturbation response scanning of gene regulatory networks

#### Preprocessing of Gene Regulatory Networks

Let *A* ∈ ℝ^*p*×*p*^ represent the adjacency matrix of the GRN, where *p* is the number of genes. First, the diagonal elements of *A* are set to zero to remove self-loops from the graph. Next, the network is filtered by applying an interaction cutoff, *ϕ*_*c*_, to filter out low-scoring edges. Specifically, we zero out any matrix elements *A*_*ij*_ ≤ *ϕ*_*c*_. Genes with no remaining connections (unlinked nodes) are removed from the analysis.

#### Giant component extraction and graph Laplacian construction

The filtered matrix *A* was then converted to an undirected graph *G* = (*V, E*), where *V* is the set of nodes, and *E* represents edges weighted by their interaction scores. Following the method described by Acar *et al*. [27] for modeling elastic networks and perturbation-scanning in gene networks, we then compute the largest connected component *G*_*c*_ of *G* and use this subgraph for subsequent analyses. The Graph Laplacian matrix, Γ, represents the overall connectivity of the graph.

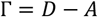

Where *D* is the degree matrix and *A* is the adjacency matrix. The diagonal elements of Γ are the degree of each node, and the non-zero, off-diagonal elements have the negative weighted interactions for respective node-pairs.

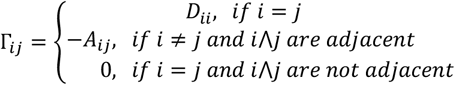

In this study, we used Γ as a medium to predict perturbations rather than focusing on pairwise interactions. Γ was then modeled as a Gaussian network model (GNM) as described by Li et al [52].

### Gaussian network modeling

The previously constructed Laplacian matrix was modeled as a GNM, where the network structure is represented as nodes connected by springs. The *n* × *n* Laplacian matrix Γ describes the dynamics and inter-node interaction topology for a network of *n* nodes.

The *n* − 1 mode spectra of the GNM were obtained through eigen decomposition of Γ, retaining only the non-zero modes. Modes were ranked by increasing frequency, with the slowest mode (mode 1) representing the first non-zero eigenpair. In GNM modelling, the eigenvalues are representative of the frequencies of the individual modes. As shown by Doruker et al. [53], in a GNM, the *k* slowest modes are crucial for understanding functional motions due to their cooperative behavior. Previous studies have shown that the collection of slowest modes reveals the mechanisms of global motions within the network [54, 55].

Hence, for the pseudoinverse computation, we performed eigen decomposition of Γ to compute the lowest *k* non-zero eigenvalues *λ*_*i*_, *i* ∈ *Z* | 1 ≤ *i* ≤ *k* and their corresponding eigenvectors *V*_*i*_, *i* ∈ *Z* | 1 ≤ *i* ≤ *k*.

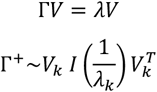

Where *λ*_*k*_ are the lowest *k* non-zero eigenvalues, *V*_*k*_ is the matrix of their corresponding eigenvectors and *I* is the identity matrix, and Γ^+^ is the approximate pseudo-inverse of Γ. The computed pseudoinverse, Γ^+^, captures the cross-correlation interactions of nodes, aiming to capture node relationship fluctuations [52].

We then constructed a Covariance matrix, *Cov*, defined as the squared elements of Γ^+^.

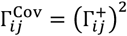

In this work, we are not directly interested in the cross-correlations captured in the Γ^Cov^ matrix; however, we will be using it to map the response of other nodes when a target node is perturbed.

#### Perturbation response matrix calculation

Perturbation response scanning (PRS) [22] was performed following one of its applications in genetic networks [27]. PRS was created for anisotropic network models (ANM) that define protein dynamics in a 3D space using Linear response theory [48]. Linear response theory for ANMs states that the total positional displacement of all nodes in the system under equilibrium is dictated by the following force balance:

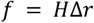

Where Δ*r* and *f* are the set of displacement and external force vectors of each node. *H* is the Hessian matrix, which is a 3*n* × 3*n* force constant matrix. This Hessian matrix represents the 3D covariance matrix of the equilibrium fluctuations of network nodes defined for protein dynamics. In the case of GNMs, this Hessian matrix is replaced by a *n* × *n* Kirchhoff matrix, also called the Laplacian matrix, Γ multiplied by a spring constant, *γ*.

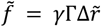

Therefore, using the Laplacian matrix, we can estimate the displacements of nodes when under an external force, *f*, using,

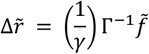

This equation can then be rewritten for the displacement of each node Δ*r*|_*i*_ in response to a force *f*_*i*_ applied on node *i* as,

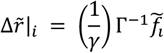

Where 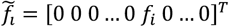, is the force exerted on the node *i*. 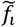 is composed of all zeros, except for the *i*^*th*^ element that is equal to one [52]. The PRS matrix of the GNM is computed by repeating the above operation for each node *i, i* ∈ (1, *N*) in the model. As the determinant of Γ is zero, the real inverse of Γ doesn’t exist. Thus, 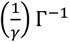 acts as a transfer function and can be replaced with the previously defined Covariance matrix, Γ^Cov^, that scales with the pseudoinverse of Γ. Therefore, the resulting PRS matrix, *δ*_*PRS*_, is

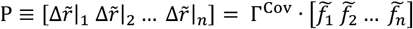

Where 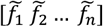 is an *n* × *n* matrix where the *i*^*th*^ column is 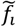, and Γ^Cov^ is the covariance matrix. The *i*^*th*^ diagonal element of Γ^Cov^ represents the square fluctuations of node *i* under equilibrium and is derived from the intrinsic property of the GNM. To ensure the perturbation responses across nodes are unitary and comparable, the response to unit deformation at each perturbation site is obtained by dividing each row by its diagonal value

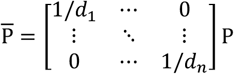

The *i*^*th*^ row represents the *influence profile* of each node that is generated upon its perturbation. The row-wise average of 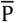 yields each node’s perturbation effectiveness, *E*. On the other hand, the column-wise average of 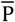, yields each node’s sensitivity to perturbations, *S*. Finally, every element 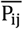 in 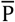 represents the response of the node *j* to the perturbation of the node *i*.

#### Gene impact score computation

Next, we use the node-specific effectiveness and sensitivity scores to quantify the overall impact of individual nodes (genes) within a gene network. The effectiveness and sensitivity scores were *log*(*x* + 1)-transformed to compress dynamic ranges and mitigate outlier effects. We apply this concept of sensor and effector nodes, characterized by structural perturbations in protein dynamics [52], to gene networks by analyzing their response to genetic perturbations, including but not limited to gene knockouts and knock-ins. Sensor nodes are characterized by their strong susceptibility to perturbations associated with their neighbor nodes. Effector nodes are characterized by the ability to effectively communicate perturbation ‘information’ to other nodes. They are located near sensor nodes, yet in tightly packed regions to minimize the dissipation of overall structural damage of the network [52].

Impact, 𝕀, was computed as the difference between effectiveness (𝔼) and sensitivity (𝕊). 𝕀 was then scaled to their respective maxima and then to the range [0,1].

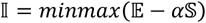

Where *α* is a tuning parameter controlling the effect of Sensitivity in the Impact computation. This was performed with the intuition that nodes with higher sensitivity are less crucial to the network topology, as they are easily influenced by changes to the network. This index provides a balanced measure of a gene’s overall impact within the network.

### Summary of network centrality measures

In addition to the proposed impact index, we computed 4 other centrality measures for each gene in the network: degree, closeness, eigenvector, and information centrality. The degree of each gene was computed directly from the Laplacian matrix:

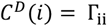

Closeness centrality, a measure of a node’s efficiency in disseminating information, was calculated as the inverse of the sum of its shortest path distances to all other nodes in the network.

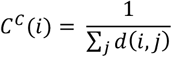

Where *C*^*C*^(*i*) is the closeness centrality of the node *i* and *d*(*i, j*) represents the shortest path between node *i* and every other node *j*.

Eigenvector centrality quantifies a node’s influence by considering both the number and ‘quality’ of its connections, such that a few highly connected and influential neighbors can confer greater importance than numerous less significant ones.

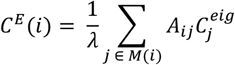

Where *C*^*E*^ (*i*) is the eigenvector centrality of the node *i. A*_*ij*_ is the element of the adjacency matrix *A*, that corresponds to the edge between nodes *i* and *j. M*(*i*) is the set of all neighbors of the node *i*, and *λ*, is the largest eigenvalue of *A*. As *A* is an undirected nonnegative adjacency matrix, by Perron-Frobenius theorem, its eigen decomposition produces a nonnegative largest eigenvector.

Information centrality, also known as, current-flow closeness centrality [56], was used to measures the extent to which a node is involved in information paths within the network.

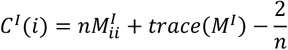

Where *C*^*I*^(*i*) is the information centrality of the node *i* and *n* is the number of vertices. *M*^*I*^ = (Γ + *J*)^− 1^ with Laplacian Γ and *J* = 11^*T*^.

All the centrality measures were computed from the graph adjacency matrix using the ‘Networkx’ Python package.

### Visualization

All graph layouts in the article were generated using the ‘spring_layout’ function in the Networkx Python package with the random seed set to 0. The enrichment bar plot was generated using the GSEApy v1.1.9 [57]. All other plots were generated using Seaborn [58].

### Hierarchical clustering and stability evaluation

To assess the biological transferability and stability of the Impact index, we obtained cell line metadata and tissue lineages from the DepMap database as our ground truth. We hypothesized that cell lines originating from similar tissues would exhibit similar centrality scores. To test this, we performed hierarchical clustering on two sets of vectors containing gene impact scores and degree centralities of all genes commonly expressed in the CCLE scRNA-seq data used in this study. Pairwise cell line distances were computed using Euclidean distance, and hierarchical linkage was performed with Ward**’**s method.

The resulting dendrograms were cut to get nine clusters, corresponding to the nine tissue types present in our data. Clustering performance for each metric was then quantitatively compared to the known tissue lineages using the Adjusted Rand Index (ARI) and Normalized Mutual Information (NMI) using the **‘**scikit-learn**’** Python library.

### Benchmarking against virtual gene KO tools

To rigorously benchmark scPII, we employed two established virtual gene KO tools: GenKI [19] and scTenifoldKnk [18]. These tools were used iteratively to perturb individual genes and quantify their systemic impact on other genes, providing a robust framework for assessing scPII’s performance. For consistency and efficiency across all three methods, we utilized pre-computed GRNs generated by GRNBoost2.

For GenKI, scRNA-seq data were processed and fed into a variational graph autoencoder (VGAE), as previously described by Yang et al. [19]. This VGAE learned latent representations of genes and their interactions from the wild-type (WT) scRNA-seq data. In each iterative step, a single gene was designated as the target for virtual KO. The effect of each gene’s KO was then quantified by calculating the distance between the latent space representations of the KO and WT conditions. Specifically, the Kullback-Leibler (KL) divergence served as a measure of dissimilarity between the two conditions in the learned latent space. These individual gene KO responses were stored in a response matrix where rows represent the KO effect on genes and columns represent the knocked-out genes, with values indicating the KL divergence of the VGAE outputs. Diagonal elements, representing a gene’s effect on itself, were set to zero to provide a standardized baseline for comparison.

For scTenifoldKnk, the same iterative single-gene KO strategy was employed. Each gene’s KO was simulated by setting its corresponding row in the GRN adjacency matrix to zero, as described by Osorio et al. [18]. The resulting perturbed GRN was then aligned to the wild-type (WT) GRN using manifold alignment. The Euclidean distances between gene projections in the low-dimensional manifold served as a quantitative measure of the global response induced by each gene’s perturbation. These gene KO responses were compiled into a response matrix, where rows denote the effect on specific genes, and columns represent the knocked-out genes, with diagonal elements (self-perturbation effects) set to zero for standardized comparison.

All benchmarking and evaluations were performed on a dedicated machine equipped with a 14-core Intel Core i5-13500 CPU processor clocked at 2.50 GHz and 31.7 GB of available RAM.

### Gene set enrichment analysis

Enrichment analysis was performed using the GSEAPY Python package, querying the DSigDB database [31] via Enrichr [30] with default settings to identify enriched drugs. Enriched terms were considered significant at an adjusted P value of less than 0.05.

## Code Availability

scPII is publicly available as a Python package, along with all codes used to generate results in this study, at https://github.com/Xenon8778/scPII.

## Data Availability

scRNA-seq data of CCLE cancer cell lines were obtained from the Broad Institute’s single-cell portal (SCP542). CRISPR screens for cancer cell lines were obtained from DepMap Public 24Q2 Release. The Glioblastoma project network was obtained from BioGRID release 4.4.243, and associated CRISPR KO screens were obtained from BioGRID-ORCS release 2.0.18. No new data was generated in support of this research.

## Acknowledgements

This research was funded by the Cancer Prevention and Research Institute of Texas (CPRIT, RP230204) and the U.S. Department of Defense (DoD, GW200026), and the National Institute for Environmental Health Sciences (P30 ES029067) for J.J.C.

## Author information

### Contributions

S.G.: Conceptualization, Methodology, Visualization, Software, Formal Analysis, Writing - Original Draft. J.J.C: Conceptualization, Methodology, Supervision, Writing - Original Draft, Resources, Software. S.R.: Writing - Review & Editing, Software.

## References

[1] A.-L. Barabási, Z. N. Oltvai, A.-L. Barabási, and Z. N. Oltvai, “Network biology: understanding the cell’s functional organization,” Nature Reviews Genetics 2004 5:2, vol. 5, no. 2, 2004/02, doi: 10.1038/nrg1272.

[2] S. G. Hofmann, “A Network Control Theory of Dynamic Systems Approach to Personalize Therapy,” Behavior Therapy, vol. 56, no. 1, 2025/01/01, doi: 10.1016/j.beth.2024.10.006.

[3] Z. M et al., “Current and future directions in network biology - PubMed,” Bioinformatics advances, vol. 4, no. 1, 08/14/2024, doi: 10.1093/bioadv/vbae099.

[4] S. Manicka et al., “The nonlinearity of regulation in biological networks,” npj Systems Biology and Applications 2023 9:1, vol. 9, no. 1, 2023-04-04, doi: 10.1038/s41540-023-00273-w.

[5] L. Wang et al., “Dictys: dynamic gene regulatory network dissects developmental continuum with single-cell multiomics,” Nature Methods 2023 20:9, vol. 20, no. 9, 2023-08-03, doi: 10.1038/s41592-023-01971-3.

[6] R. Edwards and L. Glass, “Combinatorial explosion in model gene networks,” Chaos: An Interdisciplinary Journal of Nonlinear Science, vol. 10, no. 3, 2000/09/01, doi: 10.1063/1.1286997.

[7] E. Kim et al., “A network of human functional gene interactions from knockout fitness screens in cancer cells,” Life Science Alliance, vol. 2, no. 2, 2019 Apr 12, doi: 10.26508/lsa.201800278.

[8] F. Markowetz, “How to Understand the Cell by Breaking It: Network Analysis of Gene Perturbation Screens,” PLoS Computational Biology, vol. 6, no. 2, 2010 Feb 26, doi: 10.1371/journal.pcbi.1000655.

[9] A. Dixit et al., “Perturb-Seq: Dissecting Molecular Circuits with Scalable Single-Cell RNA Profiling of Pooled Genetic Screens,” Cell, vol. 167, no. 7, 2016/12/15, doi: 10.1016/j.cell.2016.11.038.

[10] G. S and C. JJ, “Gene function revealed at the moment of sitochastic gene silencing,” Communications biology, vol. 8, no. 1, 01/19/2025, doi: 10.1038/s42003-025-07530-0.

[11] M. F. Rasul et al., “Strategies to overcome the main challenges of the use of CRISPR/Cas9 as a replacement for cancer therapy,” Molecular Cancer 2022 21:1, vol. 21, no. 1, 2022-03-03, doi: 10.1186/s12943-021-01487-4.

[12] D. Kim et al., “Gene regulatory network reconstruction: harnessing the power of single-cell multi-omic data,” npj Systems Biology and Applications 2023 9:1, vol. 9, no. 1, 2023-10-19, doi: 10.1038/s41540-023-00312-6.

[13] M. Stock, N. Popp, J. Fiorentino, and A. Scialdone, “Topological benchmarking of algorithms to infer gene regulatory networks from single-cell RNA-seq data,” Bioinformatics, vol. 40, no. 5, 2024/05/02, doi: 10.1093/bioinformatics/btae267.

[14] A. Pratapa et al., “Benchmarking algorithms for gene regulatory network inference from single-cell transcriptomic data,” Nature Methods 2020 17:2, vol. 17, no. 2, 2020-01-06, doi: 10.1038/s41592-019-0690-6.

[15] M. Zhao, W. He, J. Tang, Q. Zou, and F. Guo, “A comprehensive overview and critical evaluation of gene regulatory network inference technologies,” Briefings in Bioinformatics, vol. 22, no. 5, 2021/09/02, doi: 10.1093/bib/bbab009.

[16] T. Moerman et al., “GRNBoost2 and Arboreto: efficient and scalable inference of gene regulatory networks,” Bioinformatics, vol. 35, no. 12, 2019/06/15, doi: 10.1093/bioinformatics/bty916.

[17] J. Gillis and P. Pavlidis, “The role of indirect connections in gene networks in predicting function,” Bioinformatics, vol. 27, no. 13, 2011 May 6, doi: 10.1093/bioinformatics/btr288.

[18] D. Osorio et al., “scTenifoldKnk: An efficient virtual knockout tool for gene function predictions via single-cell gene regulatory network perturbation,” Patterns, vol. 3, no. 3, 2022/03/11, doi: 10.1016/j.patter.2022.100434.

[19] Y. Yang et al., “Gene knockout inference with variational graph autoencoder learning single-cell gene regulatory networks,” Nucleic Acids Research, vol. 51, no. 13, 2023/07/21, doi: 10.1093/nar/gkad450.

[20] C. Chennubhotla and I. Bahar, “Signal Propagation in Proteins and Relation to Equilibrium Fluctuations,” PLOS Computational Biology, vol. 3, no. 9, Sep 21, 2007, doi: 10.1371/journal.pcbi.0030172.

[21] R. Bonetto, H. J. Kojakhmetov, R. Bonetto, H. J. Kojakhmetov, R. Bonetto, and H. J. Kojakhmetov, “Nonlinear diffusion on networks: Perturbations and consensus dynamics,” Networks and Heterogeneous Media 2024 3:1344, vol. 19, no. 3, 2024, doi: 10.3934/nhm.2024058.

[22] C. Atilgan and A. R. Atilgan, “Perturbation-Response Scanning Reveals Ligand Entry-Exit Mechanisms of Ferric Binding Protein,” PLOS Computational Biology, vol. 5, no. 10, Oct 23, 2009, doi: 10.1371/journal.pcbi.1000544.

[23] S. C, B. BJ, R. T, B. L, B. A, and T. M, “BioGRID: a general repository for interaction datasets - PubMed,” Nucleic acids research, vol. 34, no.Database issue, 01/01/2006, doi: 10.1093/nar/gkj109.

[24] G. S. Kinker et al., “Pan-cancer single-cell RNA-seq identifies recurring programs of cellular heterogeneity,” Nature Genetics 2020 52:11, vol. 52, no. 11, 2020-10-30, doi: 10.1038/s41588-020-00726-6.

[25] A. Tsherniak et al., “Defining a Cancer Dependency Map,” Cell, vol. 170, no. 3, 2017/07/27, doi: 10.1016/j.cell.2017.06.010.

[26] R. Oughtred et al., “The BioGRID database: A comprehensive biomedical resource of curated protein, genetic, and chemical interactions,” Protein Science, vol. 30, no. 1, 2021/01/01, doi: 10.1002/pro.3978.

[27] O. Acar, S. Zhang, I. Bahar, and A.-R. Carvunis, “Elastic network modeling of cellular networks unveils sensor and effector genes that control information flow,” PLOS Computational Biology, vol. 18, no. 5, May 31, 2022, doi: 10.1371/journal.pcbi.1010181.

[28] L. Parkes et al., “A network control theory pipeline for studying the dynamics of the structural connectome,” Nature Protocols 2024 19:12, vol. 19, no. 12, 2024-07-29, doi: 10.1038/s41596-024-01023-w.

[29] R. Arafeh et al., “The present and future of the Cancer Dependency Map,” Nature Reviews Cancer 2024 25:1, vol. 25, no. 1, 2024-10-28, doi: 10.1038/s41568-024-00763-x.

[30] M. V. Kuleshov et al., “Enrichr: a comprehensive gene set enrichment analysis web server 2016 update,” Nucleic Acids Research, vol. 44, no. W1, 2016/07/08, doi: 10.1093/nar/gkw377.

[31] Y. M et al., “DSigDB: drug signatures database for gene set analysis - PubMed,” Bioinformatics (Oxford, England), vol. 31, no. 18, 09/15/2015, doi: 10.1093/bioinformatics/btv313.

[32] T. Eda et al., “Novel Repositioning Therapy for Drug-Resistant Glioblastoma: In Vivo Validation Study of Clindamycin Treatment Targeting the mTOR Pathway and Combination Therapy with Temozolomide,” Cancers 2022, Vol. 14, Page 770, vol. 14, no. 3, 2022-02-02, doi: 10.3390/cancers14030770.

[33] S. Berthier et al., “Anticancer properties of sodium selenite in human glioblastoma cell cluster spheroids,” Journal of Trace Elements in Medicine and Biology, vol. 44, 2017/12/01, doi: 10.1016/j.jtemb.2017.04.012.

[34] E. Yakubov, T. Eibl, A. Hammer, M. Holtmannspötter, N. Savaskan, and H.-H. Steiner, “Therapeutic Potential of Selenium in Glioblastoma,” Frontiers in Neuroscience, vol. 15, 2021 May 28, doi: 10.3389/fnins.2021.666679.

[35] M. AbdEl-haq, A. Kumar, F.-e. A. Mohand, N. Kravchenko-Balasha, Y. Rottenberg, and A. J. Domb, “Paclitaxel Delivery to the Brain for Glioblastoma Treatment,” International Journal of Molecular Sciences, vol. 24, no. 14, 2023 Jul 21, doi: 10.3390/ijms241411722.

[36] J. Lu et al., “Mechanism of action of paclitaxel for treating glioblastoma based on single-cell RNA sequencing data and network pharmacology,” Frontiers in Pharmacology, vol. 13, 2022 Nov 21, doi: 10.3389/fphar.2022.1076958.

[37] E. Macia et al., “Chlortetracycline, a Novel Arf Inhibitor That Decreases the Arf6-Dependent Invasive Properties of Breast Cancer Cells,” Molecules, vol. 26, no. 4, 2021 Feb 12, doi: 10.3390/molecules26040969.

[38] H. Tsuiki et al., “Mechanism of hyperploid cell formation induced by microtubule inhibiting drug in glioma cell lines,” Oncogene 2001 20:4, vol. 20, no. 4, 2001-03-12, doi: 10.1038/sj.onc.1204126.

[39] Q. Xie et al., “Multi-omics analysis identifies glioblastoma dependency on H3K9me3 methyltransferase activity,” NPJ Precision Oncology, vol. 9, no. 1, 2025 Mar 20, doi: 10.1038/s41698-025-00829-5.

[40] F. Lefranc, P. Yeaton, J. Brotchi, and R. Kiss, “Cimetidine, an unexpected anti-tumor agent, and its potential for the treatment of glioblastoma (review),” (in eng), Int J Oncol, vol. 28, no. 5, pp. 1021–30, May 2006.

[41] L. F et al., “Combined cimetidine and temozolomide, compared with temozolomide alone: significant increases in survival in nude mice bearing U373 human glioblastoma multiforme orthotopic xenografts - PubMed,” Journal of neurosurgery, vol. 102, no. 4, 2005 Apr, doi: 10.3171/jns.2005.102.4.0706.

[42] G. R, Y. T, and X. W, “Enemies or weapons in hands: investigational anti-diabetic drug glibenclamide and cancer risk - PubMed,” Expert opinion on investigational drugs, vol. 26, no. 7, 2017 Jul, doi: 10.1080/13543784.2017.1333104.

[43] X. Zhang et al., “Ambroxol enhances anti-cancer effect of microtubule-stabilizing drug to lung carcinoma through blocking autophagic flux in lysosome-dependent way,” (in eng), Am J Cancer Res, vol. 7, no. 12, pp. 2406–2421, 2017.

[44] H. M et al., “Dihydroergotamine mesylate enhances the anti-tumor effect of sorafenib in liver cancer cells - PubMed,” Biochemical pharmacology, vol. 211, 2023 May, doi: 10.1016/j.bcp.2023.115538.

[45] H. DR, T. CS, H. T, and R. E, “Risk of Glioblastoma Multiforme in Patients Taking Ion Channel Blockers - PubMed,” Cureus, vol. 14, no. 10, 10/13/2022, doi: 10.7759/cureus.30277.

[46] Y. Lu et al., “Perturbation response scanning of drug-target networks: Drug repurposing for multiple sclerosis,” Journal of Pharmaceutical Analysis, vol. 15, no. 6, 2025/06/01, doi: 10.1016/j.jpha.2025.101295.

[47] S. Powers, M. DeJongh, A. A. Best, and N. L. Tintle, “Cautions about the reliability of pairwise gene correlations based on expression data,” Frontiers in Microbiology, vol. 6, 2015 Jun 26, doi: 10.3389/fmicb.2015.00650.

[48] S. Zhang et al., “ProDy 2.0: increased scale and scope after 10 years of protein dynamics modelling with Python,” Bioinformatics, vol. 37, no. 20, 2021/10/25, doi: 10.1093/bioinformatics/btab187.

[49] F. A. Wolf, P. Angerer, F. J. Theis, F. A. Wolf, P. Angerer, and F. J. Theis, “SCANPY: large-scale single-cell gene expression data analysis,” Genome Biology 2018 19:1, vol. 19, no. 1, 2018-02-06, doi: 10.1186/s13059-017-1382-0.

[50] D. Osorio, Y. Zhong, G. Li, J. Z. Huang, and J. J. Cai, “scTenifoldNet: A Machine Learning Workflow for Constructing and Comparing Transcriptome-wide Gene Regulatory Networks from Single-Cell Data,” Patterns, vol. 1, no. 9, 2020/12/12, doi: 10.1016/j.patter.2020.100139.

[51] Y. Y et al., “scTenifoldXct: A semi-supervised method for predicting cell-cell interactions and mapping cellular communication graphs - PubMed,” Cell systems, vol. 14, no. 4, 04/19/2023, doi: 10.1016/j.cels.2023.01.004.

[52] H. Li, Y.-Y. Chang, J. Y. Lee, I. Bahar, and L.-W. Yang, “DynOmics: dynamics of structural proteome and beyond,” Nucleic Acids Research, vol. 45, no. W1, 2017/07/03, doi: 10.1093/nar/gkx385.

[53] P. Doruker, A. Atilgan, and I. Bahar, “Dynamics of proteins predicted by molecular dynamics simulations and analytical approaches: Application to α-amylase inhibitor,” Proteins: Structure, Function, and Bioinformatics, 2000, doi: 10.1002/1097-0134(20000815)40:3%3C512::AID-PROT180%3E3.0.CO;2-M.

[54] B. I, E. B, J. RL, A. AR, and C. DG, “Collective motions in HIV-1 reverse transcriptase: examination of flexibility and enzyme function - PubMed,” Journal of molecular biology, vol. 285, no. 3, 01/22/1999, doi: 10.1006/jmbi.1998.2371.

[55] L. Yang, G. Song, and R. L. Jernigan, “How Well Can We Understand Large-Scale Protein Motions Using Normal Modes of Elastic Network Models?,” Biophysical Journal, vol. 93, no. 3, 2007 May 4, doi: 10.1529/biophysj.106.095927.

[56] U. Brandes and D. Fleischer, “Centrality Measures Based on Current Flow,” STACS 2005, 2005, doi: 10.1007/978-3-540-31856-9_44.

[57] Z. Fang, X. Liu, and G. Peltz, “GSEApy: a comprehensive package for performing gene set enrichment analysis in Python,” Bioinformatics, vol. 39, no. 1, 2023/01/01, doi: 10.1093/bioinformatics/btac757.

[58] M. L. Waskom, “seaborn: statistical data visualization,” Journal of Open Source Software, vol. 6, no. 60, 2021/04/06, doi: 10.21105/joss.03021.

