## Supplementary Figures for "Mapping Gene Impact on Single-cell Transcriptomic Networks via Perturbation Response Scanning"

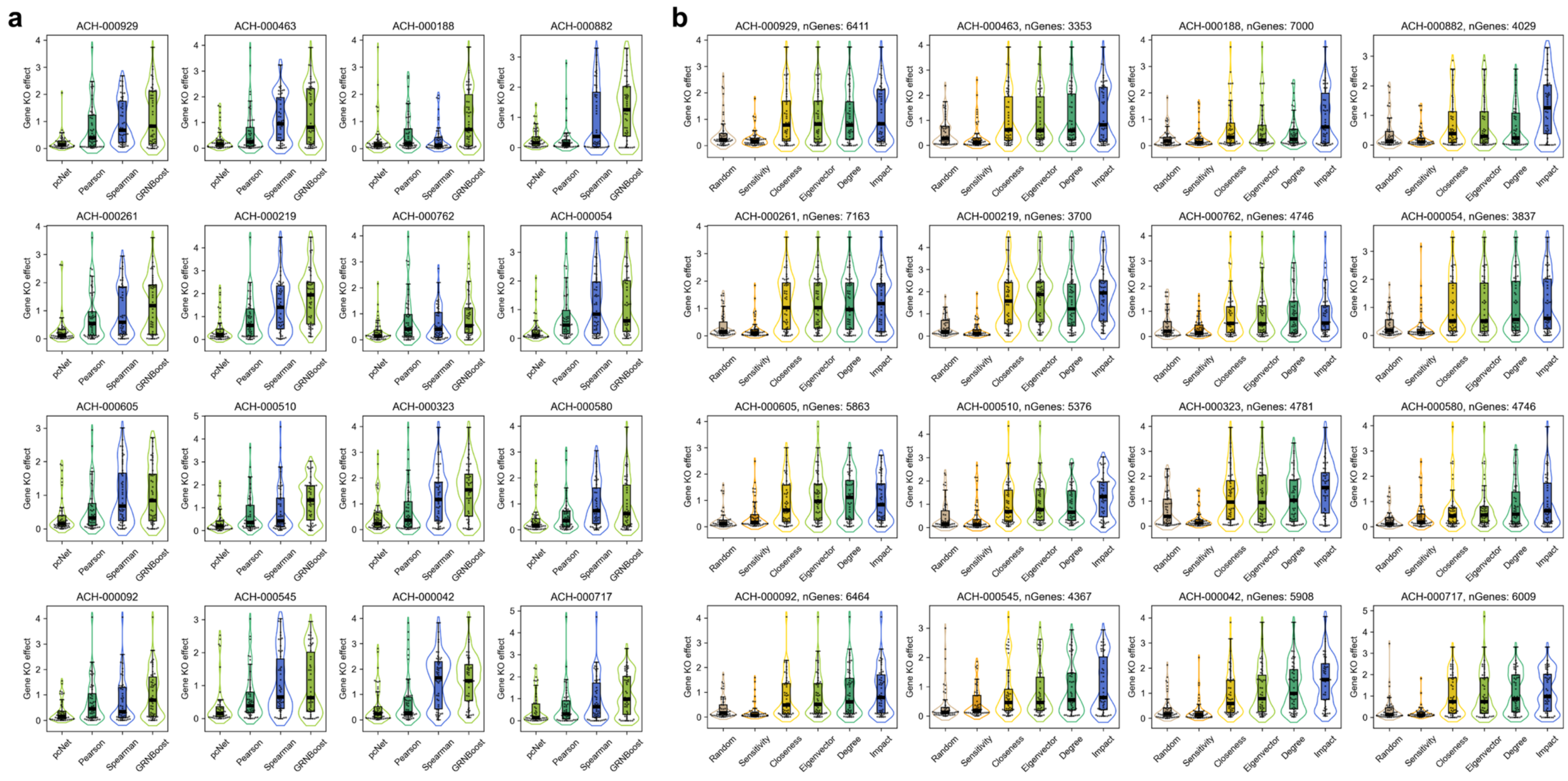


**Supplementary Figure 1.** Violin and boxplots showing gene KO effect of top 50 genes from CCLE datasets according to **(a)** Impact metric for respective GRN methods and **(b)** respective centrality measures from GRNBoost2-derived GRN.

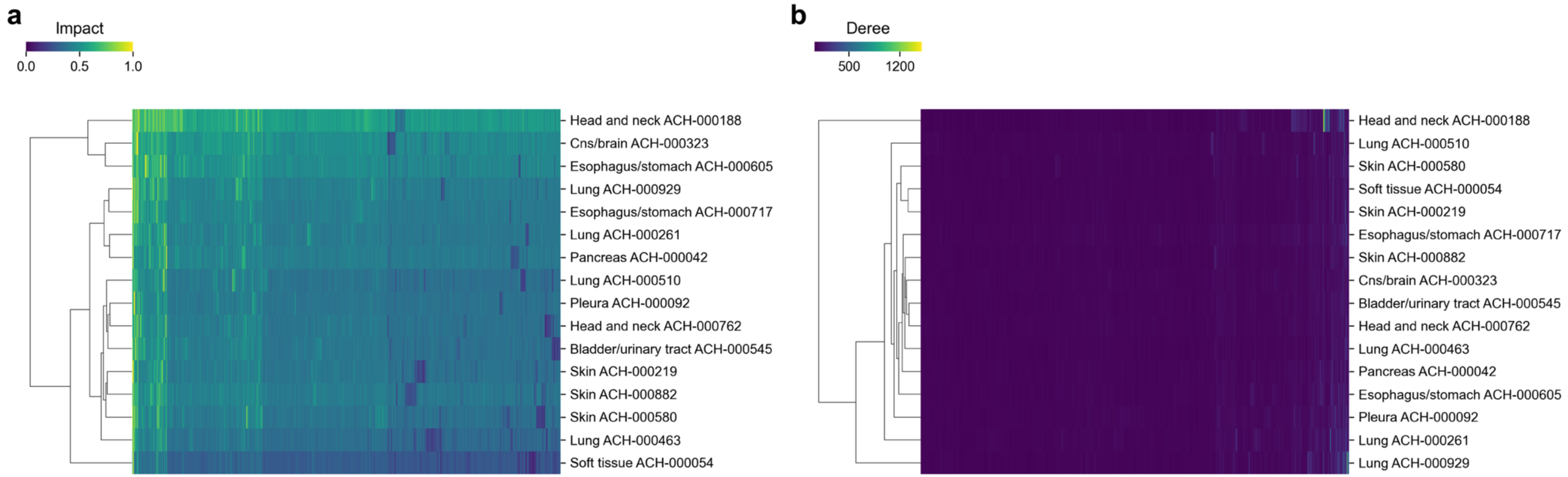


**Supplementary Figure 2.** Heatmaps showing clusters of CCLE cell lines according to **(a)** Impact and **(b)** Degree centrality scores.
